# Genomic surveillance framework and global population structure for *Klebsiella pneumoniae*

**DOI:** 10.1101/2020.12.14.422303

**Authors:** Margaret M. C. Lam, Ryan R. Wick, Stephen C. Watts, Louise T. Cerdeira, Kelly L. Wyres, Kathryn E. Holt

## Abstract

*K. pneumoniae* is a leading cause of antimicrobial-resistant (AMR) healthcare-associated infections, neonatal sepsis and community-acquired liver abscess, and is associated with chronic intestinal diseases. Its diversity and complex population structure pose challenges for analysis and interpretation of *K. pneumoniae* genome data. Here we introduce Kleborate, a tool for analysing genomes of *K. pneumoniae* and its associated species complex, which consolidates interrogation of key features of proven clinical importance. Kleborate provides a framework to support genomic surveillance and epidemiology in research, clinical and public health settings. To demonstrate its utility we apply Kleborate to analyse publicly available *Klebsiella* genomes, including clinical isolates from a pan-European study of carbapenemase-producing *Klebsiella*, highlighting global trends in AMR and virulence as examples of what could be achieved by applying this genomic framework within more systematic genomic surveillance efforts. We also demonstrate the application of Kleborate to detect and type *K. pneumoniae* from gut metagenomes.

## TEXT

*Klebsiella pneumoniae* bacteria commonly colonize the mammalian gut, but are also recognized as a major public health threat due to their ability to cause severe infections in healthcare settings and their association with antimicrobial resistance (AMR)^1,2^. Reports of *K. pneumoniae* gut colonization frequencies vary by country and demographics, but prevalence rates as high as 87% have been reported^3–6^. *K. pneumoniae* colonization is implicated in chronic diseases of the gastrointestinal tract including inflammatory bowel disease and colorectal cancer^7^. There is also a growing body of evidence highlighting colonization as a reservoir for extraintestinal infections (urinary tract infection, pneumonia, wound or surgical site infections, sepsis) in vulnerable individuals such as neonates, the elderly, immunocompromized and hospitalized patients^8^. Treatment of healthcare-associated (HA) *K. pneumoniae* infections is often limited by multidrug resistance (MDR) resulting from the accumulation of horizontally-acquired AMR genes and mutations in core genes^2^. Treatment is further complicated by increasing frequencies of strains producing extended-spectrum β-lactamases (ESBL) and/or carbapenemases, prompting increased reliance on colistin and β-lactamase inhibitor combinations^9,10^. The World Health Organization has accordingly prioritized *K. pneumoniae* as a target for new drugs and therapies^11^.

Outside healthcare settings, *K. pneumoniae* is also recognized as a causative agent of community-acquired infections including urinary tract infection and pneumonia, but also invasive infections such as pyogenic liver abscess, endophthalmitis and meningitis^12,13^. Invasive community-acquired infections are generally associated with so-called hypervirulent *K. pneumoniae* (hvKp) and are most commonly reported in East and Southeast Asia, or in individuals with East Asian ancestry^12^. Features associated with hvKp include a K1, K2 or K5 polysaccharide capsule and horizontally-acquired virulence factors encoding the siderophores aerobactin (Iuc) and salmochelin (Iro), the genotoxin colibactin (Clb), and a hypermucoid phenotype (conferred by the *rmpADC* locus)^14–18^. HvKp are rarely MDR and most remain susceptible to drugs except ampicillin, to which *K. pneumoniae* are intrinsically resistant due to the chromosomally-encoded β-lactamase SHV^19^. However there have been increasing reports of hvKp carrying AMR plasmids and co-occurrence of AMR and virulence determinants in non-hvKp isolates. The convergence of AMR and virulence in *K. pneumoniae* potentiates invasive and difficult-to-treat infections, and at least one fatal outbreak has been documented in China where carbapenemase-producing hvKp are increasingly common^20–24^.

Research conducted in the pre-genomic era characterized 77 distinct capsular (K) serotypes^25^, nine O types^26^ and variable AMR profiles amongst the *K. pneumoniae* population^27,28^, indicating a diverse genetic and phenotypic landscape^15,29^. In recent years, genomic studies have provided key insights into the population structure of *K. pneumoniae* (recently summarized in Wyres et al^16^), revealing hundreds of deep-branching phylogenetic lineages comprising sequence types (STs) or clonal groups (CGs) defined by the seven-gene multi-locus sequence typing (MLST) scheme^29^. Some of these lineages correspond to lineages (i.e. STs and CGs) that have accumulated large numbers of AMR genes that have become globally distributed (e.g. CG258, CG15, ST307); these are dubbed MDR clones and have been linked with HA infections and hospital outbreaks worldwide^30^. Others carry a high load of virulence genes (e.g. CG23, CG65, CG86) and are recognized as hvKp associated with community-acquired infections. Further distinguishing MDR from hvKp clones are their K and O antigen profiles, with the former displaying a diverse range of K and O biosynthesis loci as a result of homologous recombination between strains, while hvKp rarely deviate from the K1, K2 or K5 types^16^.

Importantly, genomic characterization of clinical isolates identified as *K. pneumoniae* via biochemical tests or mass spectrometry (MALDI-TOF) has revealed the existence of multiple related species and subspecies, which together form the *K. pneumoniae* species complex (KpSC). These differ by 3-4% nucleotide divergence across core chromosomal genes, but share the same pool of AMR and virulence genes^16^. Infections and outbreaks caused by other KpSC members have been reported but they generally account for a significantly lower disease burden than *K. pneumoniae* (10-20%)^19,31,32^. Genomics has also clarified that the two *K. pneumoniae* subspecies originally defined by distinct and unusual disease manifestations (subsp. *rhinoscleromatis* which causes a progressive and chronic granulomatous infection known as rhinoscleroma, and subsp. *ozaenae* which causes atrophic rhinitis or ozena) actually represent CGs of *K. pneumoniae* (CG3 and CG90)^15^. Like hvKp clones, these strains also express specific capsule types (K3, K4 and K5) alongside aerobactin and another acquired siderophore, yersiniabactin (Ybt)^16^.

Due to its clinical importance and increasing AMR, *K. pneumoniae* is increasingly the focus of surveillance efforts and molecular epidemiology studies. The sheer volume of clinically-relevant molecular targets renders whole genome sequencing (WGS) the most cost-efficient characterization approach, however extracting and interpreting clinically important features is challenging. To address this, we have developed Kleborate, a genotyping tool designed specifically for *K. pneumoniae* and the associated species complex, which consolidates detection and genotyping of key virulence and AMR loci alongside species, lineage (ST) and predicted K and O antigen serotypes directly from genome assemblies. Here we describe Kleborate and demonstrate its utility by application to publicly available datasets. First, we show that Kleborate can rapidly recapitulate and augment the key findings from a recent large-scale European genomic surveillance study^33^. Next, we apply Kleborate to a curated collection of 13,156 publicly available WGS to further showcase its utility and derive novel insights into the global epidemiology of *Klebsiella* AMR, virulence and convergence. Finally, we show that Kleborate can also be applied to detect clinically relevant genotypes from meta-genome assembled genomes (MAGs).

## RESULTS

### Integrated genomic framework and genotyping tool

Our goal was to develop a single tool that can rapidly extract genotype information that is clinically relevant to *K. pneumoniae* and other members of the species complex in order to support genomic epidemiology and surveillance. We have previously reported genotyping schemes for the acquired *K. pneumoniae* virulence loci *ybt, clb, iuc* and *iro*^34,35^ (whose detection and typing is implemented in early versions of Kleborate), and also K and O antigen typing implemented in the software Kaptive^36^. Here we expand the Kleborate framework to include additional features including taxonomic assignment to species and subspecies, assignment to lineages via seven-locus MLST, detection and genotyping of the *rmp* hypermucoidy locus and the *rmpA2* gene, and identification of AMR determinants (mutations and horizontally acquired genes, including assignment of SHV β-lactamase alleles as either ESBL, β-lactamase inhibitor resistance, or intrinsic ampicillin resistance only, see **Methods** and **Supplementary text)**. Kleborate can optionally call Kaptive for K/O antigen prediction.

Unlike generic AMR or virulence typing tools, we include only genetic features for which there is strong evidence of an associated phenotype in *K. pneumoniae* that has confirmed clinical relevance. These are reported in a manner that facilitates interpretation, including summarizing virulence and AMR genotypes into scores that reflect escalating clinical risk in *K. pneumoniae* infections. Kleborate features are summarized in **Table 1** and methodological details for genotyping are provided in **Methods**. For a typical 5.5 Mbp genome, a Kleborate run including AMR typing takes <10 seconds on a laptop, while robust K and O serotype prediction using Kaptive^36^ adds an additional ~1 minute. Results are output in tab-delimited format, making it easy to integrate Kleborate into existing workflows.

**Table 1.** Genome features reported by Kleborate.

| Feature | Description |
| --- | --- |
| Assembly quality | Contig count, N50, largest contig, ambiguous bases |
| Identification | Species<sup>16</sup>, MLST<sup>29,67</sup> (if *K. pneumoniae* species complex) |
| Acquired virulence determinants | Presence, genotypes, associated MGEs, truncations<br>• yersiniabactin<sup>34</sup>,<br>• colibactin<sup>34</sup>,<br>• aerobactin<sup>35</sup>,<br>• salmochelin<sup>35</sup>,<br>• hypermucoidy loci *rmpADC* and *rmpA2* |
| Virulence score | 0=no yersinabactin, colibactin or aerobactin;<br>1=yersiniabactin only; 2=yersiniabactin and colibactin (or colibactin only); 3= aerobactin without yersiniabactin or colibactin; 4= aerobactin with yersiniabactin (no colibactin); 5=yersiniabactin, colibactin and aerobactin |
| Serotype prediction | *wzi* allele and associated K locus<sup>60</sup> (default),<br>Full K and O locus typing via *Kaptive*<sup>36</sup> (optional) |
| AMR determinants (optional) |  |
| Acquired genes | Total count, alleles grouped by drug class, truncations |
| Mutations in core genes | SHV beta-lactamase (ESBL or inhibitors)<sup>68</sup>,<br>OmpK35/OmpK36 osmoporins<sup>41,42</sup> (carbapenems),<br>MgrB/PmrB<sup>69–71</sup> (colistin), GyrA/ParC<sup>72</sup><br>(fluroquinolones) |
| Number of drug classes | Excludes penicillins since resistance is intrinsic |
| Resistance score | 1=ESBL; 2=Carbapenemase; 3=Carbapenemase plus colistin resistance; 0 otherwise |

**Table 2.** Prevalence of virulence loci, ESBL and carbapenemase genes in non-redundant *Klebsiella* genomes

| Species Complex | Species | Total no. genomes | Virulence prevalence | ESBL prevalence | Carbapenemase prevalence |
| --- | --- | --- | --- | --- | --- |
| *K. pneumoniae* species complex | *K. pneumoniae* | 9705 | Ybt: 4309, 44%<br>Clb: 794, 8%<br>Iuc: 1090, 11%<br>Iro: 683, 7%<br>Rmp: 782, 8%<br>RmpA2: 716, 7% | 4634, 48% | 4173, 43% |
|  | *K. quasipneumoniae* subsp. *quasipneumoniae* | 119 | - | 31, 26% | 29, 24% |
|  | *K. quasipneumoniae* subsp. *similipneumoniae* | 363 | Ybt: 8, 2%<br>Iuc: 6, 2%<br>Iro: 4, 1%<br>Rmp: 4, 1%<br>RmpA2: 3, 0.8% | 138, 38% | 32, 9% |
|  | *K. quasivariicola* | 16 | - | 3, 19% | - |
|  | *K. africana* | 1 | - | - | - |
|  | *K. variicola* subsp. *variicola* | 498 | Ybt: 15, 3%<br>Iuc: 4, 0.8%<br>Iro: 5, 1%<br>Rmp: 4, 0.8%<br>RmpA2: 2, 0.4% | 52, 10% | 36, 7% |
|  | *K. variicola* subsp. *tropica* | 18 | - | 2, 11% | 1, 6% |
| *K. oxytoca* species complex | *K. oxytoca* | 98 | Ybt: 96, 98% | 9*, 9% | 6, 6% |
|  | *K. grimontii* | 75 | Ybt: 41, 55% | 1*, 1% | 3, 4% |
|  | *K. huaxiensis* | 4 | - | 1*, 25% | - |
|  | *K. michiganensis* | 144 | Ybt: 102, 71%<br>Clb: 1, 0.7% | 21*, 15% | 33, 23% |
|  | *K. pasteurii* | 21 | Ybt: 21, 100% | 1*, 5% | - |
|  | *K. spallanzanii* | 4 | - | -* | - |
| NA | *K. aerogenes* | 209 | Ybt: 101, 48%<br>Clb: 95, 45%<br>Iro: 190, 91% | 24, 11% | 34, 16% |
|  | *K. indica* | 2 | - | - | - |

\*excluding OXY genes that are conserved in *K. oxytoca* species complex

#### Species and subspecies assignment

The taxonomy of *Klebsiella* is rapidly evolving, with several new species and subspecies recently identified^37–39^. As a consequence, many genomes in public databases are incorrectly assigned. We therefore introduced a custom approach for rapid and accurate species and subspecies identification for *Klebsiella*, based on Mash distances^40^ to a taxonomically-curated genome set (representative tree in **Figure S1A-B**), avoiding the need for users to download large reference genome databases (see **Methods**). This approach was validated using a set of n=285 diverse clinical isolates and compared with species assignments based on the read-based taxonomic classifier Kraken2 (details in **Supplementary Text**, **Table S1, Figure S1**).

#### Virulence and AMR scores

Genomes are scored according to the clinical risk associated with the AMR and virulence loci that are detected (see **Methods**). Here we take advantage of the structured distribution of AMR and virulence determinants within the *K. pneumoniae* population^14^ to reduce the genotyping data to simple numerical summary scores that reflect the accumulation of loci contributing to clinically relevant AMR or hypervirulence: virulence scores range from 0 to 5, depending on the presence of key loci associated with increasing risk (yersiniabactin < colibactin < aerobactin); resistance scores range from 0 to 3, based on detection of genotypes warranting escalation of antimicrobial therapy (ESBL < carbapenemase < carbapenemase plus colistin resistance, see **Table 1**). These simple numerical scores facilitate downstream analyses including trend detection. For example, analysis of a non-redundant subset of 9,705 publicly available *K. pneumoniae* genomes (see below, **Table S2**) showed increasing AMR and virulence scores over time (barplots in **Figure 1A-B**). The virulence and resistance scores were correlated not only with the prevalence of individual components that contribute to the scores, but also with other components that are co-distributed in the population (lines in **Figure 1A-B**). For example, the frequencies of *rmpADC* and *rmpA2* loci over time were correlated with the virulence score (**Figure 1A**); and the resistance score was correlated with the mean number of acquired AMR genes and associated drug classes (excluding ESBLs, carbapenemases and colistin which contribute to the score) (**Figure 1C**). Consistent with this, genomes with resistance scores >0 (assigned based on the presence of ESBL and/or carbapenemase genes) typically carry many additional AMR genes conferring resistance to multiple drug classes (**Figure 1D-E**). Reducing the data to key axes of virulence and AMR also facilitates exploration of subpopulations associated with AMR, virulence or convergence of both traits; such as specific *K. pneumoniae* lineages or specimen types (see below).

**Figure 1.**
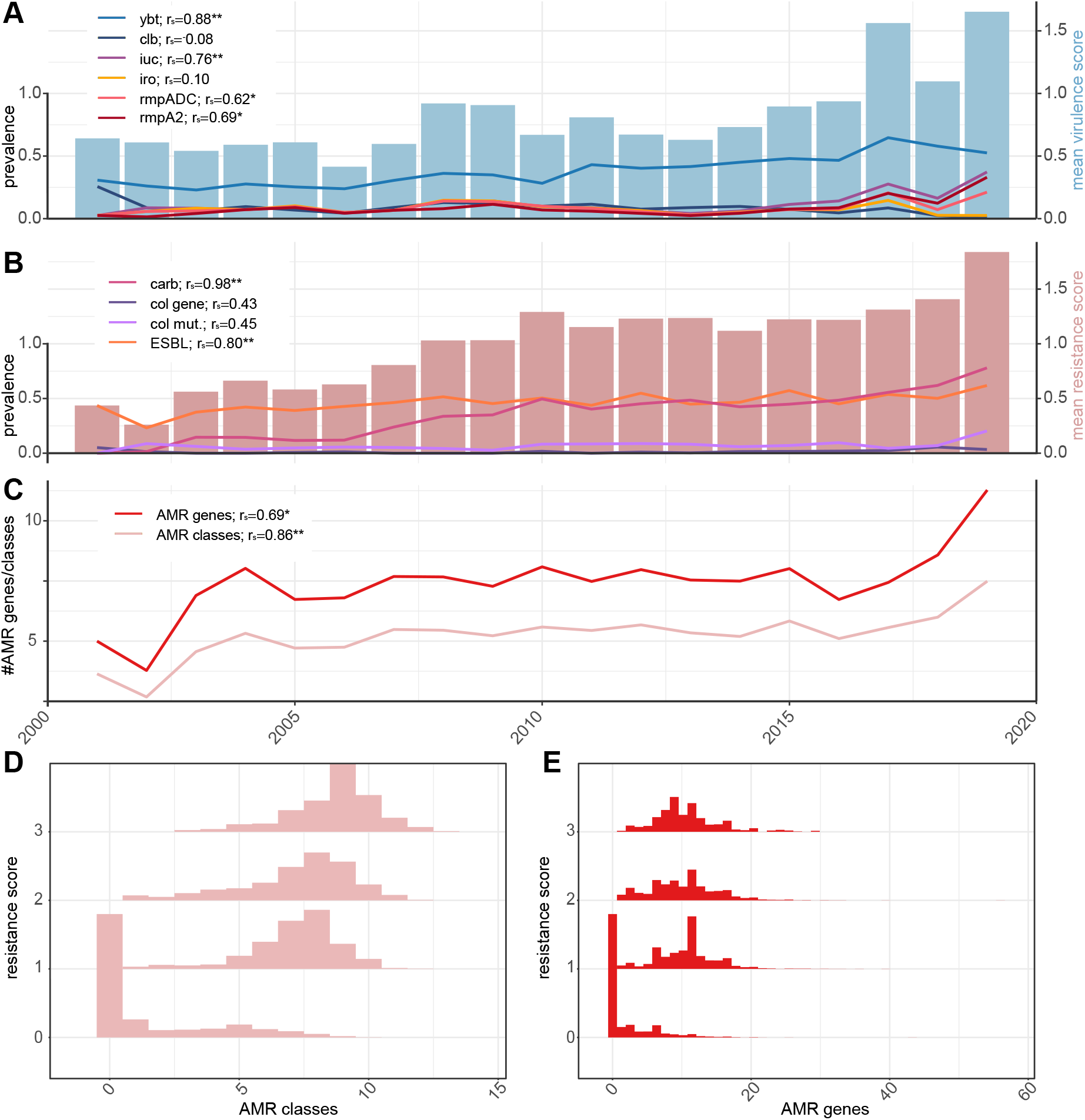
Relationships between Kleborate virulence and resistance scores and the prevalence of key virulence and antimicrobial resistance (AMR). Data shown summarise Kleborate results for non-redundant set of 9,705 publicly available *K. pneumoniae* genomes (**Table S2**). **(A)** Mean virulence score (barplot, right y-axis) and prevalence of individual virulence loci (line plots, left y-axis) over time. Ybt, yersiniabactin; clb, colibactin; iuc, aerobactin; iro, salmochelin; rmpADC, hypermucoidy *rmp* locus; rmpA2, *rmpA2* gene. Correlations between mean virulence score and prevalence of each locus are noted. **(B)** Mean resistance score (barplot, right y-axis) and prevalence of carbapenemases (carb), acquired colistin resistance genes (col gene), mutations in MgrB/PmrB (col mut) and genes conferring resistance to extended-spectrum β-lactams (ESBL) (line plots, left y-axis). Correlations between mean resistance score and prevalence of each resistance type are noted. **(C)** Mean number of acquired AMR genes and classes, over time. Correlations with mean resistance score are noted. **(D)** Histograms showing total number of acquired AMR classes predicted per genome, stratified by resistance score. **(E)** Histograms showing total number of acquired AMR genes detected per genome, stratified by resistance score. Correlations reported in **A-C** are Spearman rank correlations; significance levels are indicated with asterisks: *p<0.01, **p<0.001.

### Rapid genotyping of clinical isolates from a large-scale surveillance study

We applied Kleborate to analyse all *K. pneumoniae* clinical isolate genomes deposited in RefSeq by the EuSCAPE surveillance study (927 carbapenem-non-susceptible, 697 carbapenem-susceptible; see **Table S2**)^33^. Kleborate rapidly and accurately reproduced key findings from the original study, which were originally derived from multi-step analyses comprising five independent tools and four independent databases (each from a different public repository, one with additional manual curation): (i) 70.2% of carbapenem-non-susceptible genomes (n=651/927) carried carbapenemases, mainly KPC-3, OXA-48, KPC-2 and NDM-1; (ii) these were dominated by a few major clones, ST11, ST15, ST45, ST101, ST258, and ST512; (iii) individual countries were associated with specific carbapenemase/clone combinations (see **Figure 2A**). A detailed comparison of the results reported by Kleborate versus those reported in the original study is provided in **Supplementary Text** and **Table S3**.

**Figure 2.**
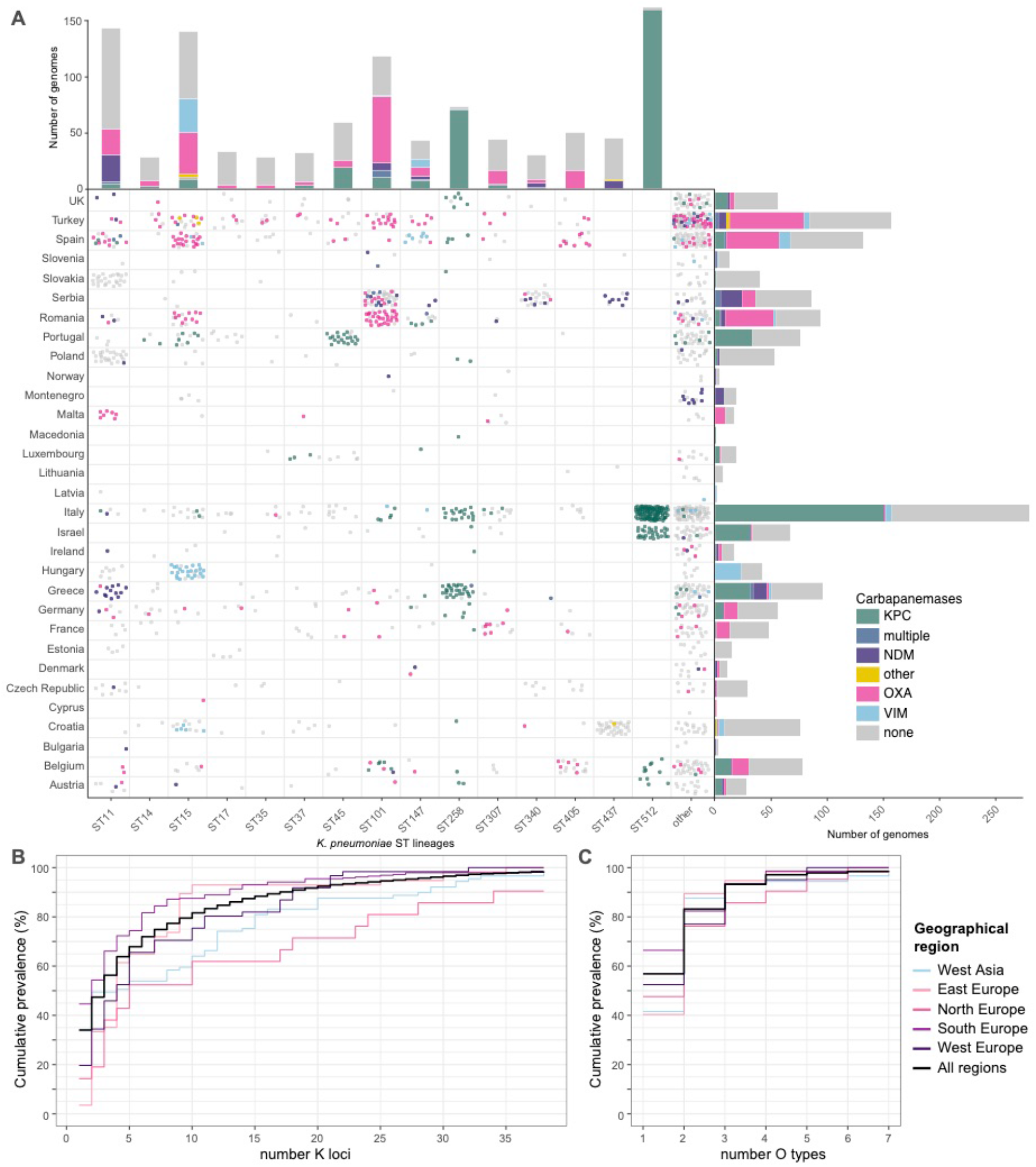
Kleborate genotyping results for European *K. pneumoniae* surveillance isolates. Data shown summarise Kleborate results for 927 carbapenem-non-susceptible and 697 carbapenem-susceptible *K. pneumoniae* genomes from the EuSCAPE study (data included in **Table S2**). **(A)** Geographical and lineage distribution of carbapenemase genes. Each circle represents a genome, colored by carbapenemase (see inset legend). Barplots summarise the number of genomes from each *K. pneumoniae* lineage (top) and country (right), colored by carbapenemase. (**B-C**) Cumulative prevalence of **(B)** capsule (K) locus and **(C)** O antigen locus types, for carbapenem non-susceptible (meropenem MIC>2) isolates, ordered by overall prevalence. Thick line indicates curve for whole data set; others give results separately for different United Nations geographical regions (see inset legend).

In addition to the detection of carbapenemase genes, Kleborate also identified porin defects, which are known to contribute to the carbapenem-resistance phenotype^41,42^, in 36.5% of EuSCAPE genomes (including 60% of those with carbapenemase genes and 19.9% of those without carbapenemase genes). These defects included truncation/deletion of OmpK35 and/or OmpK36 (also considered in the original study) as well as GD or TD insertions in the OmpK36 β-strand loop^41^ (not considered in original study, but here detected in 18.6% of genomes including 18 with no porin deletion). **Figure 3** shows meropenem MICs stratified by porin defect-carbapenemase combinations identified by Kleborate, highlighting the importance of porin defects – including the OmpK36 β-strand loop insertions – for full expression of carbapenem resistance in *K. pneumoniae*.

**Figure 3.**
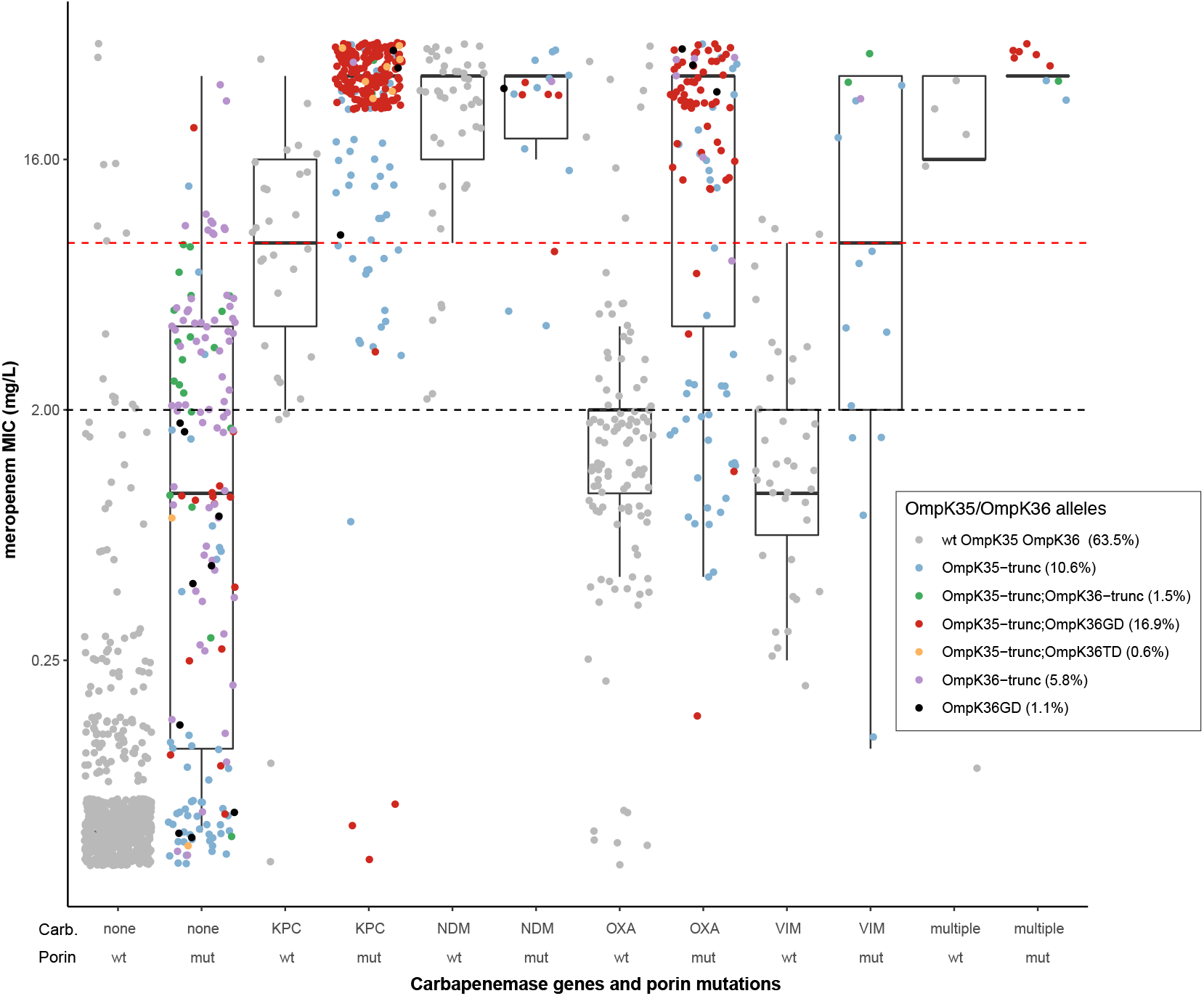
Distribution of meropenem MIC, stratified by Kleborate-detected carbapenemase genes and OmpK35/36 porin mutations, for European *K. pneumoniae* surveillance isolates. Data shown summarise Kleborate results for 1,490 *K. pneumoniae* genomes from the EuSCAPE study (data included in **Table S2**). Each circle represents the reported meropenem MIC for an isolate, coloured by type of porin mutation/s identified by Kleborate from the corresponding genome assembly (colour key in inset legend, prevalence of each genotype across 1490 genomes is indicated in brackets). Isolates are stratified by carbapenemase gene (enzymes labelled on x-axis) and OmpK mutations^41,42^ reported by Kleborate. Wt, full-length OmpK35 and OmpK36 with no GD/TD insertion in the OmpK36 β-strand loop; mut, otherwise; trunc, truncation. Dashed lines indicate EUCAST breakpoints for clinical resistance (red, MIC >8) and non-susceptibility (black, MIC >2).

The rise in carbapenem-resistant *K. pneumoniae* infections in hospitals and its associated morbidity and mortality^43^ has led to increased interest in alternative control strategies such as vaccines, phage therapy and antibody therapy, key targets for which are the K and O surface antigens^44,45^. Kleborate confidently identified K and O biosynthesis loci in 98.3% and 99.1% of EuSCAPE genomes, respectively, including 87 distinct K loci and 11 distinct O loci (**Figures S2** and **S3**). Amongst carbapenem-non-susceptible isolates (meropenem MIC >2), 38 distinct K types were identified and the most common were KL107 (n=173), KL17 (n=67), KL106 (n=41), KL24 (n=35), KL15 (n=19) and KL36 (n=13). Seven distinct O types were detected among these genomes, and the most common were O2 (n=294), O1 (n=136) and O4 (n=52). Overall, the data suggest an intervention would need to be effective against six K types or two O types in order to provide coverage of 80% of carbapenem-resistant infections in Europe (**Figure 2B-C**). However, it is important to explore the impact of population structure on these findings, specifically the impact of local clonal expansions. Kleborate aides this type of analysis by providing ST and other genotyping information alongside the K and O locus types, which can be viewed in the context of geographic information. Doing so revealed that each of the top three K loci were dominated by a single ST (83.5% of KL107 were ST512; 93.0% of KL105 were ST11; 91.4% of KL17 were ST101). Importantly, the vast majority of ST512-KL107 genomes (75.3%) originated from Italy where this ST is known to be locally circulating^46,47^, while 58% of ST11-KL105 originated from Poland and Slovakia, and 56% of ST101-KL17 originated from Serbia and Romania. When these putative local expansions were excluded, the top 6 K loci were (KL24, KL15, KL2, KL112, KL107, KL151) and accounted for just 34% of the remaining genomes.

### Global population snapshot of *K. pneumoniae* AMR and virulence

We applied Kleborate to analyse n=13,156 *Klebsiella* genomes (see **Methods**, **Table S2**). Here we provide a brief overview of the data followed by an exploration of AMR, virulence and the phenomenon of convergence, with the aim to highlight the rich information and types of inferences that can be derived from Kleborate output.

The genome data represented isolates collected from a range of sources in 99 countries between 1920–2020 (**Table S4**, although human isolates from the USA, China and UK dominated the data set accounting for n=4,702 genomes, 35.7% of total). The majority of these genomes were sourced from RefSeq, and among these Kleborate identified 1.0% (n=103/10,747) as a species other than the taxon recorded in NCBI; this is consistent with other studies and highlights the current confusion around taxonomic designations in *Klebsiella*. The most common species was *K. pneumoniae* (n=11,259, 86%); the rest comprised other KpSC species (9.4%), other members of the *K. oxytoca* species complex (3.1%) and *K. aerogenes* (1.9%) (**Figure 4, Table S4**). AMR and virulence genes were concentrated in the KpSC and particularly *K. pneumoniae* (**Figure 4, Table S5**).

**Figure 4.**
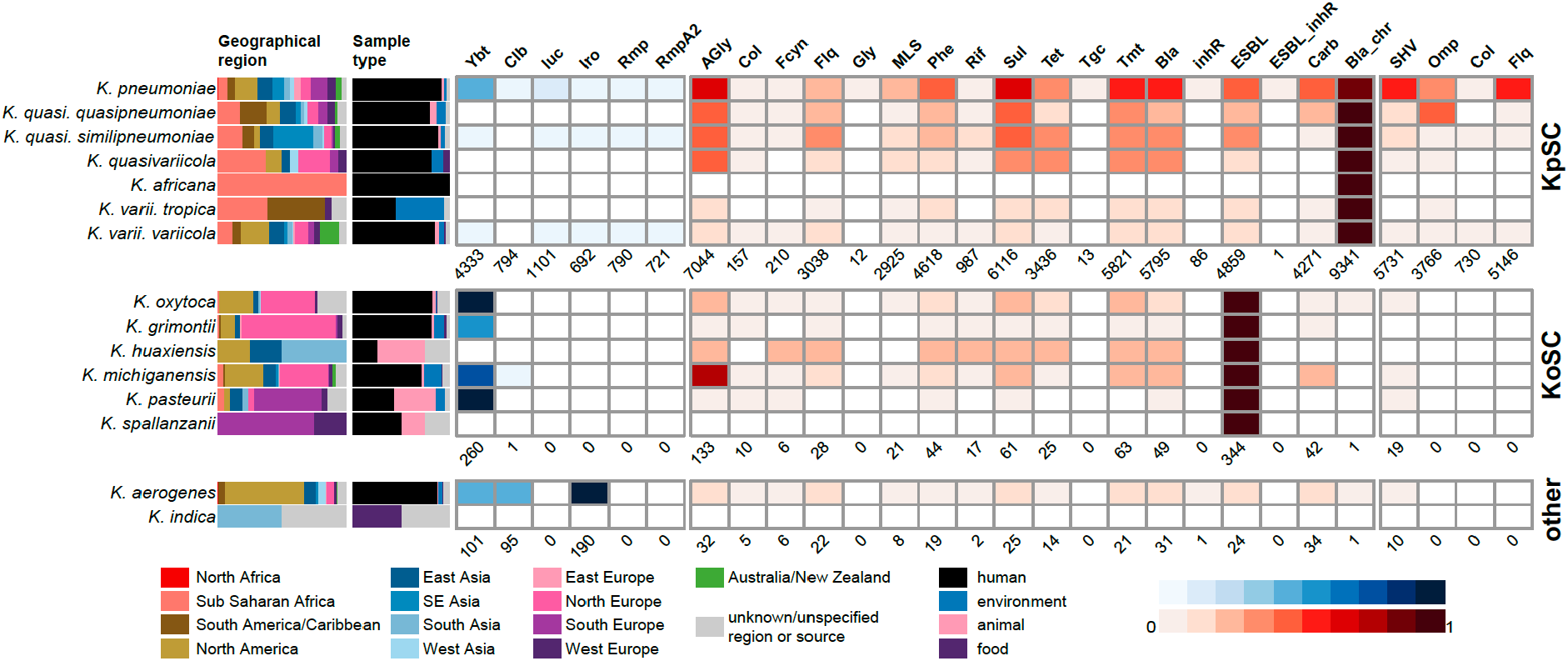
Summary of genome collection metadata, and Kleborate-derived virulence and antimicrobial resistance (AMR) genotypes, for all publicly available *Klebsiella* genomes. Data shown summarise Kleborate results for 11,277 non-redundant *Klebsiella* genomes publicly available as at July 17, 2020 (**Table S2**). From left to right: barplots showing source information by geographical region and sample type (coloured as per inset legend); heatmaps showing prevalence of virulence loci (blue) and predicted AMR drug classes (red) (as per inset scale bars). Genomes are summarised by species, ordered by species complex: KpSC, *K. pneumoniae* species complex; KoSC, *K. oxytoca* species complex; and other *Klebsiella*. In the heatmaps, the total number of genomes in which each type of virulence/AMR determinant was detected are indicated below each column. Column names are as follows: ybt, yersiniabactin; clb, colibactin; iuc, aerobactin; iro, salmochelin; rmp, hypermucoidy Rmp; rmpA2, hypermucoidy rmpA2; AGly, aminoglycosides; Col, colistin; Fcyn, fosfomycin; Flq, fluoroquinolone; Gly, glycopeptide; MLS, macrolides; Phe, phenicols; Rif, rifampin; Sul, sulfonamides; Tet, tetracyclines; Tgc, tigecycline; Tmt, trimethoprim; Bla, β-lactamases; inhR, β-lactamase inhibitor; ESBL, extended-spectrum β-lactamases; ESBL_inhR, extended-spectrum β-lactamase with resistance to β-lactamase inhibitors; Carb, carbapenemase; Bla_chr, intrinsic chromosomal β-lactamase; SHV, mutations in SHV; Omp, truncations/mutations in *ompK35/ompK36*; Col, truncations in *mgrB/pmrB* conferring colistin resistance; Flq, mutations in *gyrA/parC* conferring resistance to fluoroquinolones.

The collection captured extensive phylogenetic diversity across the *K. pneumoniae* species (see interactive phylogeny at http://microreact.org/project/JDyan46yctyDh6weEUjWN), and Kleborate assigned these genomes to ≥1,452 different STs (1,119 known STs across and at least 333 novel STs). Notably, 600 STs (41%) were represented by just a single genome each (accounting for 5.3% of all genomes). We detected n=4 ST67 (subspecies *rhinoscleromatis*) and n=3 ST90 (subspecies *ozanae*). A small number of STs were overrepresented, reflecting the bias towards sequencing MDR and hypervirulent isolates, as well as those causing hospital outbreaks. For example, 1,354 genomes (12.0%) represented the KPC-associated ST258, which is known to dominate carbapenem-resistant *K. pneumoniae* in the USA and southern Europe (where it has been the subject of intense genomic investigations) but is comparatively rare in other regions of the world^16^. To reduce the impact of these sampling biases in public genome collections, we down-sampled to a non-redundant set of 9,705 *K. pneumoniae* genomes representing unique combinations of ST, genetic subcluster (Mash distance <0.0003), virulence genotype, AMR genotype, specimen type, location and year of isolation (see **Methods**). However, we cannot fully correct for the sampling biases inherent in the public genome data and even after subsampling, the 30 most common STs accounted for 63.4% of genomes (n≥50 genomes each, n=6,151 total; see **Figure S4**). **Figure 5** shows the distribution of AMR and virulence scores amongst non-redundant genomes from these 30 common *K. pneumoniae* STs (n>50 per ST), each of which displays high rates of AMR and/or virulence.

**Figure 5.**
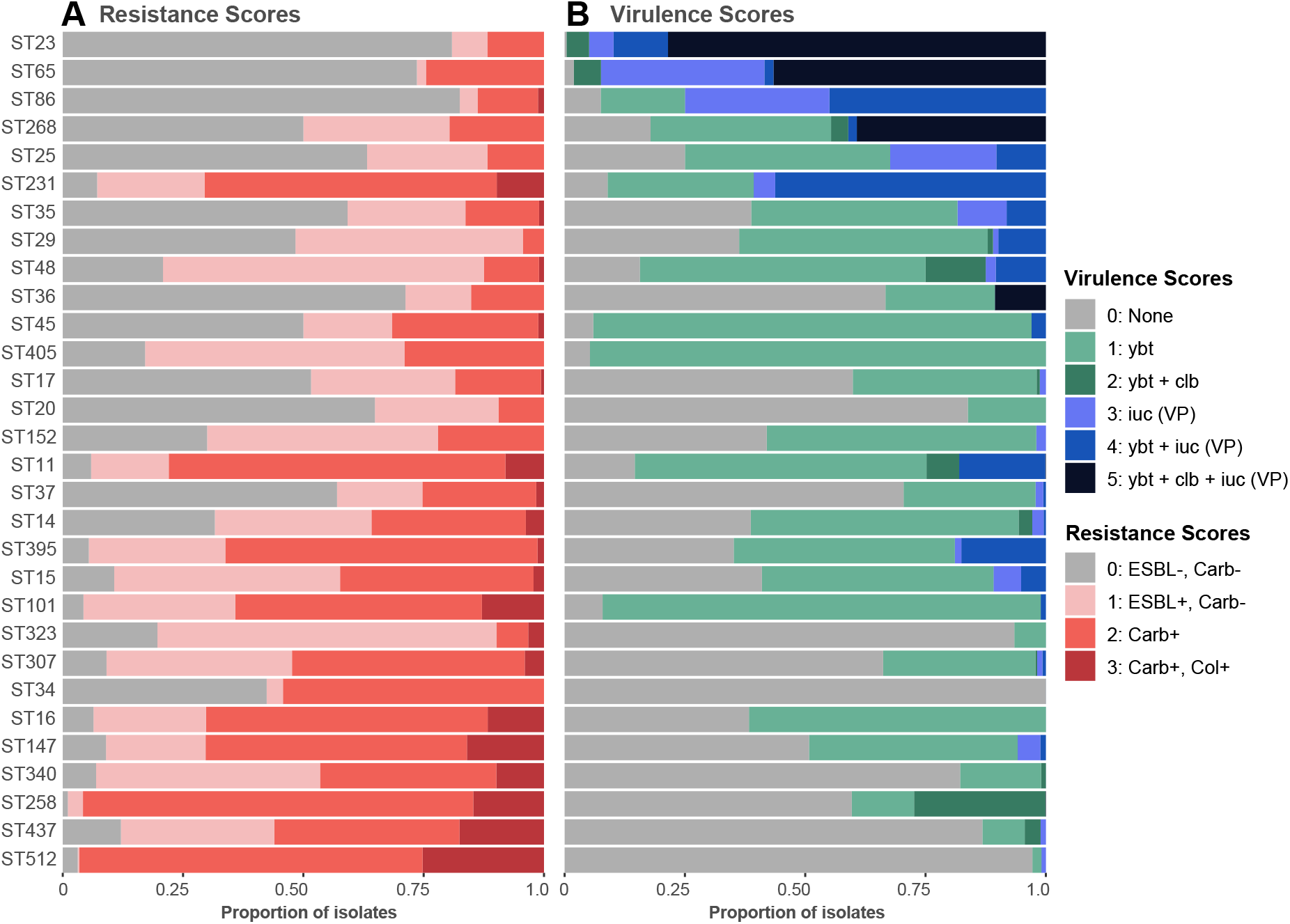
Distribution of resistance and virulence scores among genomes belonging to the 30 most common *K. pneumoniae* lineages. Data shown summarise Kleborate results for non-redundant set of 9,705 publicly available *K. pneumoniae* genomes (**Table S2**). Lineages were defined on the basis of multi-locus sequence types (STs) reported by Kleborate, and ordered from highest to lowest difference between mean virulence and mean resistance score. Minimum genome count per ST shown is 50. Ybt, yersiniabactin; clb, colibactin; iuc, aerobactin; VP, virulence plasmid; ESBL, extended-spectrum β-lactamase; Carb, carbapenemase; Col, colistin resistance determinant

#### AMR determinants

SHV β-lactamases conferring intrinsic resistance to the penicillins were detected in 85.9% of the 9,705 non-redundant *K. pneumoniae* genomes (ESBL forms of SHV were detected in 10.0%). Acquired AMR was widespread (77.1% of genomes had at least one gene or mutation conferring acquired AMR detected) and 71.6% of genomes were predicted to be MDR (acquired resistance to ≥3 drug classes^48^), a much higher rate than is reported in most geographical regions^3,49–51^, reflecting the bias within public genome collections. The majority of genomes had a non-zero resistance score, reflecting presence of ESBL and/or carbapenemase genes: 22.3%, 37.1% and 5.9% genomes had resistance scores of 1, 2 and 3 respectively. Mean resistance scores increased through time (**Figure 1B**). This trend could be an artefact of sampling bias towards the selective sequencing of AMR isolates, however it is consistent with the increasing AMR rates reported in surveillance studies globally^52–54^.

Comparatively higher prevalence of acquired AMR genes was observed in some STs (**Figure S4**). Many of these STs represent recognized MDR clones largely from clinical samples that were also associated with high mean resistance scores (**Figures 6A-B**), driven by high frequency of ESBL and carbapenemase genes (**Figures 5, S5A-B**). The most common ESBLs/carbapenemases were widely detected across the population (46-299 STs each), including amongst the top 30 common STs (prevalence range per ST, 0.1-100%; see **Figure S5A-B**), highlighting their mobile nature. The notable exception was CTX-M-65, which appeared to be largely clone specific, detected in only 9 STs and ST11 accounting for 96.7% of these genomes.

**Figure 6.**
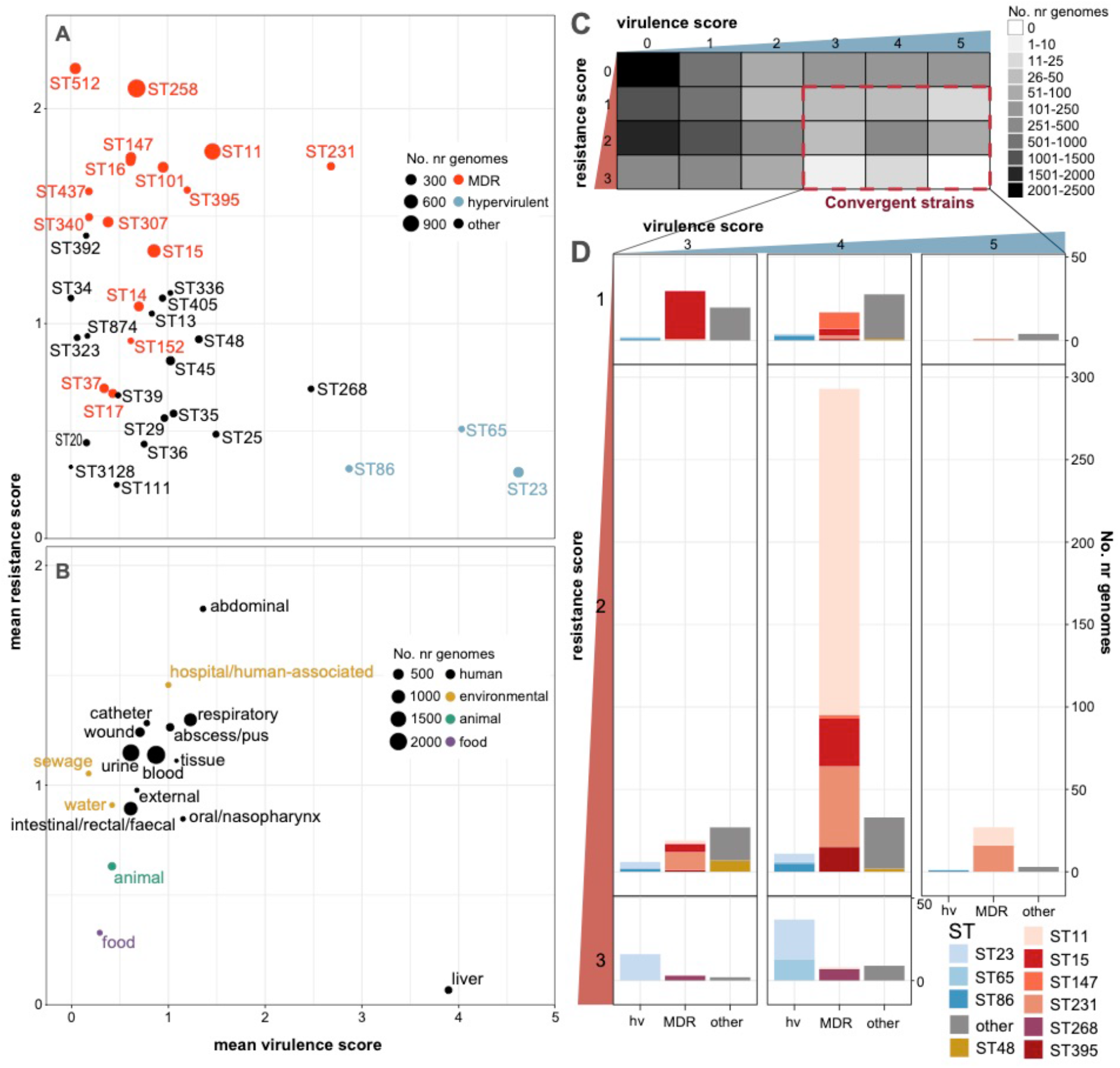
Insights from resistance and virulence scores. Data shown summarise Kleborate results for non-redundant set of 9,705 publicly available *K. pneumoniae* genomes (**Table S2**). (**A-B**) Mean resistance and virulence scores grouped by (**A**) lineage and (**B**) sample type. Each circle represents a single lineage (multi-locus sequence type, ST) or sample type as labelled; size indicates the number of genomes (as per inset legend); colour indicates groups per inset legend. **(C)** Heatmap showing number of genomes with each combination of resistance and virulence scores. Convergent genomes correspond to a virulence score ≥3 (carrying *iuc*) and resistance score of ≥1 (carrying ESBL and/or carbapenemase gene/s), as indicated by the red box. **(D)** Barplots showing lineage distribution of convergent genomes, for each combination of resistance score and virulence score. Lineages are grouped into hypervirulent (hv), multidrug resistant (MDR) and other; and coloured by ST (as per inset legend).

Colistin resistance determinants were detected in 8.7% of the non-redundant *K. pneumoniae* genomes. These were mostly nonsense mutations in MgrB or PmrB (83.5%) rather than acquisition of an *mcr* gene (15.8%, and an additional 6 genomes with both acquired *mcr* and truncated MgrB/PmrB). The rate of detection ranged from 0-25.2% for the 30 most common STs, and was highest amongst ST512, ST437, ST147, ST16 and ST258 (**Figure S5C)**, each of which are also associated with high rates of carbapenem-resistance. Porin mutations were detected in 37.9% of genomes (34.0% OmpK35, 20.2% OmpK36, 16.3% both). High prevalence of specific porin defects have been reported previously in some clones^41,42^, and this was reflected in our analysis of ST258 and its derivative ST512. We observed OmpK35 truncations in 99.9% of non-redundant ST258 genomes (with or without truncations or substitutions in OmpK36), and truncations in OmpK35 and/or OmpK36 in all ST512 (99.4% with OmpK35 truncations, 94.4% with the OmpK36GD mutation, see **Figure S5D**).

#### Virulence loci

The prevalence of acquired siderophores and colibactin loci amongst non-redundant *K. pneumoniae* genomes was 44.4% *ybt*, 7.5% *clb*, 11.2% *iuc* and 7.0% *iro*. The loci were found across diverse *K. pneumoniae* STs (391 STs with *ybt*, 56 with *clb*, 144 with *iuc*, 108 with *iro*) but were rarely detected in other *Klebsiella* species (with the exception of *ybt* among the *K. oxytoca* species complex, see **Figure 4**) indicating frequent mobilisation within *K. pneumoniae* but not between species (**Table S6**, **Figure S6**). Mean virulence scores increased through time (**Figure 1A**). **Figure 5B** shows the frequency of virulence scores in the top 30 most common STs in the non-redundant genome set. Sixteen of these common STs had ≥40% of genomes carrying the ICE*Kp*-associated *ybt* without the virulence plasmid-associated *iuc* locus (i.e. virulence score=1-2), including well known MDR clones ST258, ST11, ST14, ST15, ST101, ST147, ST152, ST395. Only the hvKp clones (ST23, ST86, ST65) and ST231 had a high frequency of *iuc* (virulence score ≥3).

In addition to detecting the presence of virulence loci, Kleborate reports on their completeness, genetic lineages and associated MGE variants, which can provide insights into their dissemination. Most of the virulence loci identified in the non-redundant *K. pneumoniae* data set (98%) matched one of the genetic lineages described previously^34,35^ (**Table S6**). **Figure S7A** shows the frequency of *iuc* lineages in *K. pneumoniae* STs with ≥20 non-redundant genomes and at least one genome harbouring *iuc*. There were four STs for which >60% genomes harboured *iuc*, and only a single *iuc* lineage was detected in each (*iuc1* in ST23, ST65, ST86; *iuc2A* in ST82), consistent with long-term persistence of a specific virulence plasmid in these well-known hypervirulent clones. In contrast, *iuc* was less frequent among other STs, several of which were associated with multiple *iuc* lineages (e.g. ST231, ST25, ST35), consistent with more recent and/or transient virulence plasmid acquisitions (mostly *iuc1*, followed by *iuc3* and *iuc5*).

Frameshift mutations (i.e. truncations) and/or incomplete loci (i.e. missing at least one gene) were detected in 10%, 28.5%, 13.6% and 17.7% of non-redundant *K. pneumoniae* genomes with *ybt*, *clb*, *iuc* and *iro* respectively (**Table S7**). While some of these may erroneously arise from contig breaks in draft genome assemblies, true truncations or missing genes may reflect a lack of function. The latter is likely true for instances where we observe conserved frameshift mutations across entire lineages, e.g. frameshift mutations were detected in *iucA* for all *iuc3*+ genomes and in *iroC* for all *iro3*+ and *iro4*+ genomes.

The hypermucoidy locus *rmpADC* was detected in 8.4% of non-redundant *K. pneumoniae* genomes (and just eight genomes of other KpSC species, **Table S6**). The majority of these genomes (67.2%, belonging to >79 STs) carried intact copies of all three genes, thus likely express the hypermucoid phenotype. Intact *rmpADC* was common in *iuc*-positive genomes of the hvKp clones ST23 and ST86, as well as MDR clones ST29 and ST101 (**Figure S7B**). Many other *iuc*-positive genomes carried *rmpADC* loci with truncated or missing genes, which likely do not confer the hypermucoid phenotype. Notably, these included hvKp clones ST65 and ST82, as well as MDR clones ST231, ST15 and ST14. The *rmpA2* gene was detected in 7.4% of non-redundant *K. pneumoniae* genomes, but was mostly present in truncated form (89.0% of *rmpA2*+ genomes) due to frameshifts within a poly-G tract^55^. The latter highlights the importance of considering not only the presence/absence of a given gene, but also whether it encodes a full-length protein, which may have important clinical implications.

### Facilitating detection of AMR-virulence convergence

AMR and virulence determinants have until recently been segregated in non-overlapping *K. pneumoniae* populations^14,19^, as clearly indicated by the distributions of AMR and virulence scores among STs (**Figures 5, 6A**). However, reports of convergent AMR-virulent strains with the potential to cause difficult-to-treat infections are increasingly common^16,56^. Kleborate facilitates rapid identification of such strains on the basis of resistance and virulence scores (convergence defined as virulence score ≥3 and resistance score ≥1, **Figure 6C**). Based on these scores, we observed a total of 601 convergent *K. pneumoniae* (510 non-redundant) with the highest proportion corresponding to a virulence score of 4 (indicative of yersiniabactin plus aerobactin/virulence plasmid detection) and resistance score of 2 (carbapenem resistance).

The majority of convergent genomes (74.5%) were concentrated within a small number of STs comprising the well-known hypervirulent (ST23, ST86, ST65) and MDR lineages (ST11, ST15, ST231 and ST147) (**Figures 6C-D, 7**). We combined the genotyping data and information from a Mash-distance-based neighbour-joining tree (http://microreact.org/project/JDyan46yctyDh6weEUjWN) to define unique convergence events (defined as unique combinations of ST, virulence and resistance determinants, and phylogenetic cluster). This identified n=173 convergence events, accounted for by either acquisition of the virulence plasmid by MDR/other clones (n=84 events; 475 genomes), or acquisition of ESBLs/carbapenemases by hypervirulent clones (n=89 events; 126 genomes) (**Figure 7**, **Table S8**).

**Figure 7.**
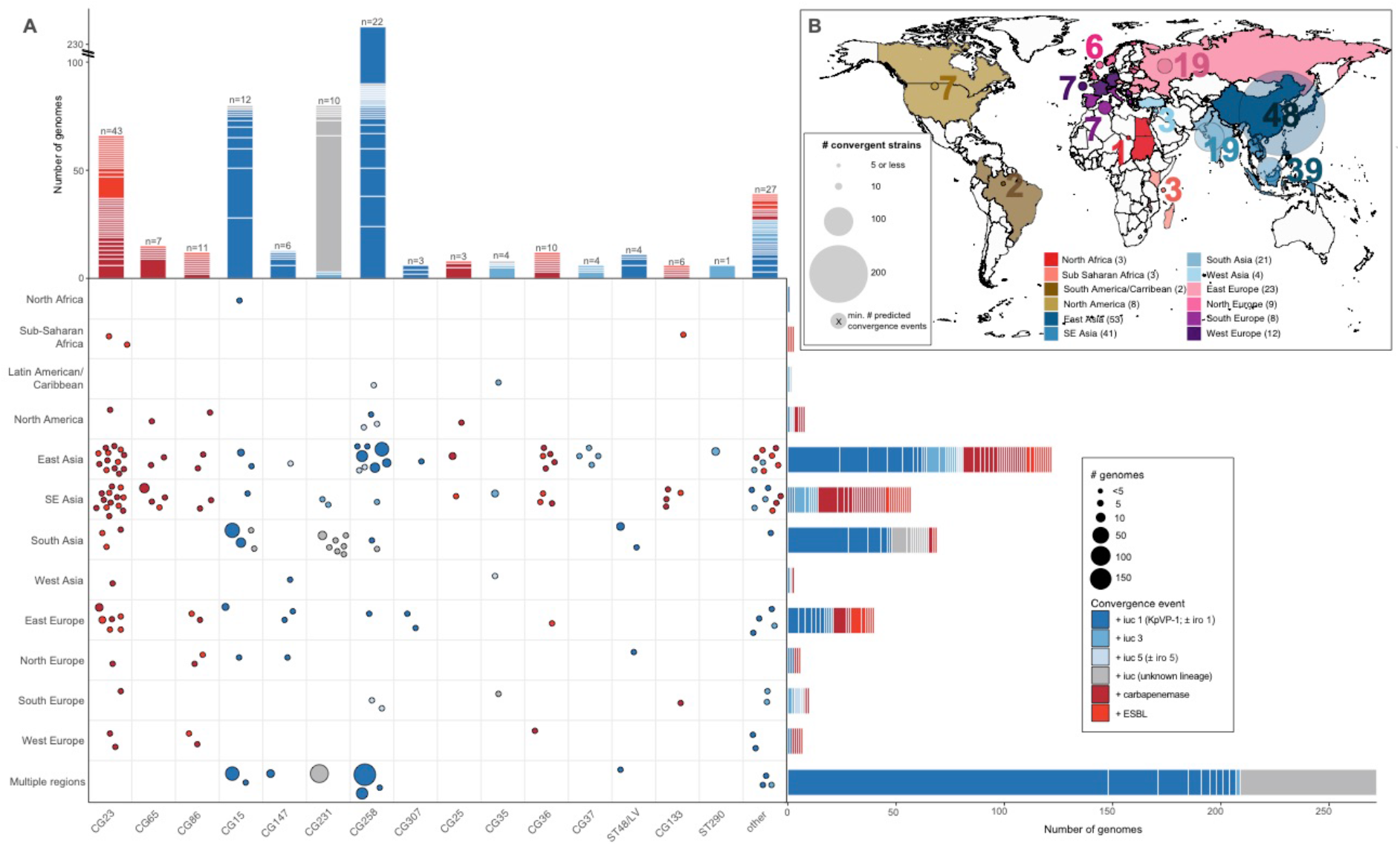
Convergence of AMR and virulence determinants in the *K. pneumoniae* population, identified by Kleborate analysis of public genomes. **(A)** Geographical and lineage distribution of convergence events. Each circle represents a unique convergence event (i.e. a monophyletic clade harbouring both ESBL/carbapenemase genes and *iuc*; see interactive tree at https://microreact.org/project/JDyan46yctyDh6weEUjWN, summary of events in **Table S8**, assignment of genomes to events in **Table S2**). Circles are scaled by the number of total genomes linked to the event and colored to indicate whether convergence is inferred to have occurred via acquisition of AMR gene/s (ESBL or carbapenemase/s) by a hypervirulent lineage or via acquisition of an *iuc*-encoding plasmid by an AMR lineage, as per inset legend. Marginal barplots show the number of convergence events (color blocks) and genomes (block heights) associated with each lineage (top) or geographical region (right). Lineages were defined on the basis of multi-locus sequence types (STs), number of convergence events estimated for each is labelled at the top of each bar. **(B)** Distribution of convergent genomes by location. Countries from which convergent genomes were detected are colored on the map; circles represent the number of convergent genomes detected in each UN-defined geographical region (indicated by color, as per inset legend), scaled and labelled with the minimum estimated number of unique convergence events specific to each region (excluding inter-regional convergence events). The total number of convergence events affecting each region, including region-specific and inter-regional convergence events, are given in brackets in the inset legend.

The most common virulence plasmid, KpVP-1 (*iuc1* ± *iro1*), accounted for 54% of virulence plasmid acquisition events (n=45 acquisitions), while *iuc3* plasmids, the *E. coli* derived *iuc5* (±*iro5*) and *iuc/iro* unknown (i.e. novel or divergent *iuc/iro* loci) accounted for 21%, 11% and 14%, respectively (**Figure 7**). AMR acquisitions by hypervirulent clones involved the ESBL/carbapenemase genes that are most common in the general *K. pneumoniae* population: KPC-2 (26%), OXA-232 (17%) and CTX-M-15 (18%). The majority of convergence events (87%) were associated with just a small number of genomes (i.e. n≤3); however, five events were associated with >20 genomes in the complete dataset, which may indicate clonal expansion and dissemination of the corresponding convergent strains locally and/or between countries. One such event corresponded to the ST11-KPC + KpVP-1 deletion variant strain that was originally reported in 2017^20^ and has since been recognized as widely distributed in China^20–24^. The complete public genome set (i.e. counting redundant genomes) included 148 genomes corresponding to this specific ST11 convergence event mostly from China but also from France (n=2). Notably though, this was only one of 50 convergence events that we detected in China, including 8 involving acquisition of *iuc1* or *iuc5* by ST11 (see **Table S8**, and interactive tree at http://microreact.org/project/JDyan46yctyDh6weEUjWN). Additional events associated with >20 genomes included (i) ST231-MDR + virulence plasmids carrying novel *iuc* lineages detected in India, Pakistan, Switzerland, Thailand and USA, (ii) ST15-CTX-M-15 + KpVP-1 in Pakistan, (iii) ST15-MDR + KpVP-1 in China and Nepal, and (iv) another distinct ST11-KPC-2 + KpVP-1 event in China. Including the above three examples, 11 convergence events appeared to involve intercountry expansion of which one has been previously documented^57^.

Overall, convergent genomes were detected originating from most geographical regions for which genome data was available, but some regions had many more events than others (**Figure 7, Table S8**). This uneven distribution may stem from a skew in the number of genomes available per region (e.g. due to variation in accessibility or application of genome sequencing). Nevertheless, the number of convergent genomes in the eastern, southeastern and southern parts of Asia were noticeably high, driven by the frequency of convergence events detected in China (n=50 events) and Thailand (n=26 events) as well as putative clonal expansions of these strains as discussed above (**Figure 7**). Of note, AMR acquisitions by hypervirulent lineages were particularly frequent within East and Southeast Asia where hypervirulent infections are most frequently reported, alongside countries from eastern and northern Europe.

Outside of *K. pneumoniae,* convergence events were rare: we detected n=2 *K. quasipneumoniae* subsp. *similipneumoniae* (ST367 with KpVP-1 and CTX-M-15; ST3387 with *iuc3* and CTX-M-55) and n=2 *K. variicola* subsp. *variicola* (ST595 with KpVP-1 and KPC-2; ST1848 with *iuc5* and KPC-2).

### Genotyping *K. pneumoniae* from metagenome data

There is increasing interest in detection and typing of *K. pneumoniae* direct from gut metagenome data^58^, due to the role of *K. pneumoniae* gut colonization as a source of acute infections and as a contributor to chronic diseases^7,8^. We tested Kleborate’s performance by application to n=40 metagenomes from which at least one KpSC isolate was cultured and sequenced, as part of the Baby Biome Study^59^. We compared the results of running Kleborate on metagenome-assembled genomes (MAGs, i.e. species-specific contig bins extracted from whole-metagenome assemblies) vs. KpSC isolate whole genome sequence(s) cultured from the same fecal sample. Thirty-two metagenomes had >1% relative abundance of KpSC, and genotyping of MAGs from these yielded results consistent with genotyping of cultured isolates for 26/32 samples (16 with identical genotypes reported for species, ST, K/O locus, virulence and AMR; 10 with close matches; see **Fig. S8, Tables S9-S10**). As expected, MAG-derived genotypes were closest to those of isolates when only one KpSC strain was cultured from the sample (see **Fig. S8, Table S10**). Kleborate analysis of whole metagenome assemblies (as opposed to individual MAGs) is not recommended: species detection and ST assignment matched that of the corresponding WGS isolates for only n=4/40 metagenome assemblies, which is unsurprising as the whole metagenomes include sequences derived from dozens of different bacteria, many of which harbour homologs of genotyping targets.

## DISCUSSION

Whole genome sequencing is being increasingly implemented in research and public health labs as a cost- and time- effective option for tracking pathogens and AMR. However, identification of clinically-relevant features remains a key bottleneck that hinders widespread adoption of genome surveillance. We have presented a comprehensive framework and tool for rapid genotyping of *Klebsiella* species genomes: Kleborate is a single unified approach for species detection, MLST and genotyping of key virulence and AMR determinants. It focusses only on genomic features for which there is strong evidence of a clinically relevant phenotype in KpSC and presents the data in a readily interpretable format, with numerical summaries and categorical scores corresponding to measures of clinical risk.

A key strength of the Kleborate framework is its species-specific approach. This is particularly important for accurate interpretation of AMR and virulence gene screens from WGS, wherein the use of generic databases and tools can result in confusion. Notable examples include the intrinsic *oqxAB* and *fosA* alleles, which unlike for other Enterobacterales, do not confer resistance to quinolones and fosfomycin when expressed in KpSC. Kleborate does not report these intrinsic alleles, neither does it report intrinsic virulence determinants such as the siderophore enterobactin, which is known to play a role in KpSC pathogenicity but for which the presence alone cannot be considered to indicate enhanced virulence of one isolate over another. Correct taxonomic identification of *K. pneumoniae* can be difficult in itself, hence the inbuilt speciation tool is an important feature (and here identified nearly 100 RefSeq genomes with incorrect species/subspecies assignments).

Another strength of our approach is the rich data output by Kleborate, which facilitates in-depth investigation of population structure, AMR and virulence epidemiology. This allows rapid exploration and understanding of: (i) hypervirulence-associated loci and the molecular drivers of their dissemination (**Figures S4** and **S7**); (ii) molecular mechanisms of complex AMR phenotypes e.g. carbapenem resistance (**Figure 3**); (iii) AMR and virulence trends (**Figures 1**, **5** and **6**); (iv) emerging convergent AMR-virulent strains so that they can be targeted for surveillance and infection control (**Figure 7**); (v) overrepresented STs and genotypes, which may be indicative of transmission clusters that should be targeted for further investigation (as demonstrated for the EuSCAPE surveillance genomes, **Figure 2A**); (vi) surface antigen epidemiology, which can inform the design of novel vaccines and therapeutics (**Figure 2B-C**). Notably Kleborate can also yield useful genotyping results from metagenomics data (**Figure S8**), which is gradually being adopted for clinical and surveillance applications relevant to *K. pneumoniae*. User interpretation of Kleborate’s extensive data output can be guided by the accompanying web-based visualization app, Kleborate-Viz. Through this app, many of the analyses and plots presented in this manuscript can be rapidly replicated, and further explored in an interactive manner.

Kleborate is designed to facilitate detection and tracking of clinically relevant AMR and virulence traits from genome data, and analysis of public data not only identified specific clones and genes associated with one or the other of these traits (**Figures 5, 6**), but also 601 genomes in which the two converge (carrying *iuc*+ virulence plasmids and ESBL and/or carbapenemase genes; **Figure 7**). We estimated at least 173 unique AMR-hypervirulence convergence events; the majority were detected within a single isolate (n=119 events), but many others appear to be associated with local outbreaks or larger-scale spread and apparently across multiple countries (**Table S8**). Some of the convergence events in China and other countries in the neighbouring South and Southeast Asia regions have been extensively reported^16,20,49,56^, but to our knowledge a significant number had not been recognized previously. These include ST231-MDR (most with OXA-232, remainder with ESBL only) + *iuc*, which has been reportedly circulating in India^49^, and our analysis also detected in Pakistan, Thailand, Switzerland and USA.

Kleborate has already been widely adopted by the *Klebsiella* research community – at least 74 studies have reported using the Kleborate software package, including larger-scale genome surveillance studies in South and Southeast Asia, the Caribbean and the United States^31,49,50^ (full list in **Table S11**). Kleborate is freely available as a standalone command-line tool for local high-throughput analyses or incorporation into existing bioinformatics workflows (https://github.com/katholt/Kleborate), and can be easily accessed through the online tool PathogenWatch (https://pathogen.watch/). With such broad accessibility and utility, Kleborate is poised to become a cornerstone of the *Klebsiella* genomic surveillance toolkit that can help inform containment and control strategies targeting this priority pathogen.

## METHODS

### Kleborate software: implementation and genotyping logic

Kleborate (v.2) is a command-line tool written in Python and is freely available under the GNU v3.0 license at http://github.com/katholt/Kleborate. It takes as input one or more whole genome assemblies (FASTA format), types each one against a series of screening databases outlined in detail below, and returns results in a tab-delimited text file (one genome per row). On default settings, Kleborate will report assembly quality metrics, taxonomic assignment, MLST and virulence loci genotypes. Screening for AMR determinants, and/or K/O serotyping via Kaptive^36^, is optional (**Table 1**).

#### Assembly quality

Assembly quality metrics, reported to help users assess the reliability of genotyping results, are: contig count, contig N50, largest contig size, total genome size, and number of ambiguous bases (e.g. ‘N’). Low quality warnings are flagged if: (i) ambiguous bases are detected; (ii) assembly length falls outside the expected range of 4.5-7.5 Mbp; or (iii) N50 is below 10,000 bp. Users should carefully consider the genotyping outputs for low quality assemblies.

#### Taxonomic assignment

Kleborate’s species prediction function provides a convenient way to confirm species, including differentiating between the closely related members of the KpSC which are frequently misclassified using laboratory techniques. Kleborate calculates Mash^40^ distances between the input genome/s and a curated collection of reference assemblies from different *Klebsiella* and other Enterobacterales, and reports the species with the smallest distance. Mash distance ≤0.02 is reported as a strong match, ≤0.04 as weak (only when no strong matches are found, see **Supplementary Text** for further details).

#### MLST

Genomes assigned to species in the KpSC are assigned sequence types using nucleotide BLAST against the established *K. pneumoniae* chromosomal seven-locus MLST scheme^29^ described and maintained on the *K. pneumoniae* BIGSdb site hosted at the Pasteur Institute (http://bigsdb.pasteur.fr/klebsiella/klebsiella.html).

#### Virulence gene detection and typing

Virulence loci (*ybt*, *iuc*, *iro*, *clb*, *rmpADC, rmpA2*) are detected using nucleotide BLAST search against the database of known alleles. The best hit allele for each gene (with ≥90% identity and ≥80% coverage) is reported in the main virulence columns. If the majority of genes expected for the locus are present, then the alleles are used to calculate STs which are reported along with their associated lineage and MGE (based on previously defined schemes: YbST for *ybt*, CbST for *clb*, AbST for *iuc*, SmST for *iro*, according to the previously defined schemes^34,35^; and a novel RmST scheme for the *rmpADC* locus). To generate the RmST typing scheme we used the same 2,733 genomes from our original virulence plasmid study^35^ to screen and extract the sequences for *rmpADC* and define allele numbers and STs. These ST sequences cluster into four distinct lineages associated with distinct MGEs (*rmp1* with KpVP-1, *rmp2* with KpVP-2, *rmp2A* with the *iuc2A* virulence plasmids, and *rmp3* with ICE*Kp1*; to be described in detail elsewhere). Where the best hit for a gene is a weak match (80-90% identity, 40-80% coverage) this is reported in the ‘spurious hits’ column. Truncations are detected by translating the best-matching nucleotide sequence for each query gene into amino acids and comparing to the reference length (expressed as % amino acid length from the start codon, those <90% are reported). The presence of *ybt*, *clb* and *iuc* are used to assign a virulence score as follows: 0=none present, 1=yersiniabactin only, 2=colibactin without aerobactin (regardless of yersiniabactin, however *ybt* is almost always present when *clb* is), 3=aerobactin only, 4=aerobactin and yersiniabactin without colibactin, and 5= all three present. The presence of *iro* (salmochelin) is not used to calculate the virulence score because its presence is very strongly associated with aerobactin.

#### Detection and typing of antimicrobial resistance determinants

When AMR detection is switched on, Kleborate screens for known acquired AMR determinants using a curated version of the CARD AMR nucleotide database (v3.0.8 downloaded February 2020; see doi.org/10.6084/m9.figshare.13256759.v1 for full details on curation). Genes are identified using nucleotide BLAST (and amino acid search with tBLASTx if no exact nucleotide match is found). Gene truncations and spurious hits are identified as described above for virulence genes. Unlike the acquired forms, the intrinsic variants of *oqxAB,* chromosomal *fosA* and *ampH* are not associated with clinical resistance in KpSC and are therefore not reported. However, SHV, LEN or OKP β-lactamase alleles intrinsic to KpSC species are known to confer clinical resistance to penicillins and are reported in the Bla_chr column. Acquired SHV variants, and individual SHV sequence mutations known to confer resistance to extended-spectrum β-lactams or β-lactamase inhibitors, are reported separately (see **Supplementary Text, Tables S12-S13** for details).

Chromosomally encoded mutations and gene loss or truncations known to be associated with AMR are reported for genomes identified as KpSC species. These include fluoroquinolone resistance mutations in GyrA (codons 83 and 87) and ParC (codons 80 and 84), and colistin resistance from truncation or loss of MgrB and PmrB (defined as <90% amino acid sequence coverage). Mutations in the OmpK35 and OmpK36 osmoporins reportedly associated with reduced susceptibility to β-lactamases^41,42^ are also screened and reported for KpSC genomes, and include truncation or loss of these genes and OmpK36GD and OmpK36TD transmembrane β-strand loop insertions^41^. SHV β-lactamase, GyrA, ParC and OmpK mutations are identified by alignment of the translated amino acid sequences against a reference using BioPython, followed by interrogation of the alignment positions of interest (see **Supplementary Text, Tables S12-S13** for a list of relevant positions).

AMR genes and mutations are reported by drug class, with β-lactamases further categorized by enzyme activity (β-lactamase, ESBL or carbapenemase, with/without resistance to β-lactamase inhibitors). Horizontally acquired AMR genes are reported separately from mutational resistance and contribute to the AMR gene count; these plus chromosomal mutations count towards the number of acquired resistance classes (intrinsic SHV alleles, reported in Bla_chr column, are not included in either count). Resistance scores are calculated as follows: 0=no ESBL or carbapenemase, 1=ESBL without carbapenemase (regardless of colistin resistance); 2=carbapenamase without colistin resistance (regardless of ESBL); 3=carbapenemase with colistin resistance (regardless of ESBL).

#### Serotype prediction

By default, genomes are screened against the *wzi* database in the *Klebsiella* BIGSdb (using nucleotide BLAST) which is used to predict capsule (K) type based on a previously defined scheme^60^. This allows rapid typing however the relationship between *wzi* allele and K type is not one-to-one^36^. If surface antigen prediction is important to users they can obtain more robust identification of K and O antigen (LPS) loci by switching on serotype prediction with Kaptive^36^ (--kaptive), which adds a few minutes per genome to Kleborate’s runtime.

#### Data visualization

To facilitate interpretation of Kleborate’s rich data output we provide a web-based application (Kleborate-Viz, https://kleborate.erc.monash.edu/), implemented in R Shiny, which takes as inputs a Kleborate results file (required), sample metadata (CSV format, optional) and MIC data (CSV format, optional).

### Genome analysis

The analyses reported here result from applying Kleborate v2.0.0 to publicly available genome collections. A total of 13,156 *Klebsiella* WGS assemblies, encompassing non-duplicate isolates with unique BioSample accessions identified from published studies (some deposited as read sets only, which were assembled using Unicycler v0.4.7^61^, data sources summarized in **Table S14**) plus any additional genomes designated as *Klebsiella* in NCBI’s RefSeq repository of genome assemblies (as of 17^th^ July 2020). In order to minimize the impact of sampling bias favouring common MDR and/or virulent lineages and those causing outbreaks, we subsampled the collection into a ‘non-redundant’ dataset of 11,277 genomes (9,705 *K. pneumoniae*) as follows. Pairwise Mash distances were calculated using Mash v2.1, and used to cluster genomes using single-linkage clustering with a threshold of 0.0003. These clusters were further divided into non-redundant groups with unique combinations of (i) Mash cluster, (ii) chromosomal ST, (iii) virulence gene profiles (i.e. presence of *ybt/clb/iro/iuc* loci and lineage assignment), (iv) AMR profiles, (v) year and country of isolation, and (vii) specimen type where available. For each resulting non-redundant group, one genome was selected at random as the representative for analyses. The full list of genomes, including database accessions, isolate information, cluster/group assignment, and Kleborate results are provided in **Table S2**. The subset of 1,624 *K. pneumoniae* assemblies deposited in RefSeq by the European EuSCAPE surveillance study^33^ (out of 1,649 reported in original study; **Table S2**) were used for the EuSCAPE analyses reported in **Figures 2** and **3**. The Kleborate-Viz web application is pre-loaded with the non-redundant and EuSCAPE WGS datasets reported in this paper, and can be used to reproduce the plots shown in **Figures 1A-C, 2B-C, 3, 6A-B** and to further explore the Kleborate results.

### Metagenome analysis

We downloaded metagenomic reads, and matched isolate WGS assemblies, for n=47 infant gut microbiota samples deposited by the Baby Biome Study^59^. Metagenome reads were assembled using SPAdes version 3.13.1^62^ with the --meta flag and the resulting contigs binned using MaxBin v2.2.7^63^. Seven metagenomes failed to assemble due to memory and compute walltime constraints, hence we report results for 40 samples (**Table S10**). Kleborate was run separately on the full metagenome assemblies, all contig bins (from which the *Klebsiella* bin could then be identified), and the matched WGS assemblies. Metagenomic read sets were also analysed using Kracken 2.0.7^64^ and Bracken v2.5^65^ (with a custom GDTB release 89 database^66^) to estimate the relative abundance of KpSC reads in each metagenome.

### Statistical analysis

Statistical analyses and data visualisations were conducted using R v1.1.456. Figures were generated with ggplot v3.2.0 and pheatmap v1.0.12. Correlations between virulence and resistance scores, and the prevalence of virulence and resistance determinants over time, were analysed using Spearman’s rank-order correlation (i.e. non-parametric test).

## Acknowledgements

We thank Prof Sylvain Brisse and the curators of the *K. pneumoniae* BIGSdb at Institut Pasteur for hosting and maintaining the MLST schemes (https://bigsdb.pasteur.fr/klebsiella/klebsiella.html); and Prof David Aanensen and team at the Centre for Genomic Pathogen Surveillance for making Kleborate available online within Pathogen<u>w</u>atch (http://pathogen.watch).

## Funding

This work was supported by the Bill and Melinda Gates Foundation of Seattle (grant OPP1175797 to K.E.H.) and the Viertel Charitable Foundation of Australia (Senior Medical Research Fellowship to K.E.H.). K.L.W is supported by the National Health and Medical Research Council of Australia (Investigator Grant APP1176192).

## Author Contributions

Study design: K.E.H. Data analysis: M.M.C.L., K.L.W. and K.E.H. Code development: R.R.W., K.E.H., S.C.W., L.T.C., and K.L.W. Manuscript writing: M.M.C.L., K.L.W., and K.E.H. All authors contributed to manuscript editing.

## Competing Interests

None declared.

## Notes

https://doi.org/10.6084/m9.figshare.c.5238239

### Competing Interest Statement

The authors have declared no competing interest.

